# Strategies for Integrating Single-Cell RNA Sequencing Results With Multiple Species

**DOI:** 10.1101/671115

**Authors:** Ronald P. Hart

## Abstract

Single-cell RNA sequencing (scRNAseq) is a robust technology for parsing gene expression in individual cells from a tissue or other complex source. One application involves experiments where cells from multiple species are recovered from a single sample, such as when human cells are transplanted into an animal model. We transplanted microglial precursor cells into newborn mouse brain and then recovered unenriched cortical tissue six months later. Dissociated cells were assessed by scRNAseq. The default method for analyzing these results begins by aligning sequencing reads with a mixture of both mouse and human reference genomes. While this clearly identifies the human cells as a distinct cluster, the clustering is artificially driven by expression from non-comparable gene identifiers from different species. We devised a method for translating expression counts from human to mouse and evaluated four algorithms for parsing mixed-species scRNAseq data. Our optimal approach split raw sequencing reads according to the best alignment score in each genome, and then re-aligned reads only with the appropriate genome. After gene symbol translation, pooled results indicate that cell types are more appropriately clustered and that differential expression analysis identifies species-specific patterns. This method should be applicable to any mixed-species scRNAseq experiment.

**Summary of optimal strategy:**

- Mixed-species scRNAseq data are aligned with mixture of mouse and human reference genomes
- The BAM file is scanned to find the best alignment score for each sequencing read identifier; these are used to split the paired FASTQ files into two sets of files
- Each set of species-specific, paired FASTQ files is re-aligned with only the appropriate reference genome
- Raw counts imported into Seurat
- The human counts table is translated to mouse gene symbols using a custom HomoloGene translation table
- Results are merged and analyzed

## Introduction

As single-cell RNA sequencing (scRNAseq) becomes widely utilized, one obstacle is the analysis of cells in a single assay from multiple species. An example application that is novel to this technology is when cells from one species are transplanted into another, and then both host and graft cells are recovered by scRNAseq for parsing into cell-level data. In our case, human iPSC-derived microglial precursor cells were implanted into newborn mouse brain (Xu et al., 2019). We recovered cortical tissue from mouse after six months of development to assess microglial gene expression patterns in both the xenograft human cells as well as the host mouse microglia. A primary goal was to compare human and mouse microglia residing in the same host brains.

In general, scRNAseq requires dissociation of tissue into single cells, followed by microfluidic packaging of individual cells into a droplet along with a single bead encoded with a barcoded primer (Macosko et al., 2015).

After cell lysis and reverse transcription from the barcoded primer within the droplet, the resulting cDNAs can be pooled and prepared for sequencing as with a traditional RNA sequencing library. We used the 10X Genomics version of the technology, with a microfluidic chamber to encapsulate cells and a design to produce paired-end sequencing results with one end (R2) giving cDNA sequence and the other (R1) revealing the cell-specific barcode sequence. The data are generally aligned with a reference genome (only the R2 reads) and counts of observed transcripts summed by barcode. An excellent review of scRNAseq technologies and analysis pipelines is available elsewhere (Hwang et al., 2018). Our goal was to process a dataset from two species. In another example, mouse and human brain scRNAseq data have been integrated for a combined cell-type analysis (Johnson et al., 2019), but this is a different design.

In our experiments, the first step in processing the samples was to align raw, barcoded sequencing results (FASTQ files) with a mixture of human and mouse reference genomes. It was assumed that each cell, identified by a unique barcode, would predominantly match only one of the two genomes. A standard workflow would then normalize/scale the data, calculate principal components, and then visualize clusters of cells in a t-distributed stochastic neighbor embedding (tSNE) plot. If only one cell type in the experiment derives from the transplanted human cells, only one tSNE cluster would be identified by expression of human genes. We observed this result as expected. However, this result poses a problem, since all other clusters are identified using mouse gene identifiers and therefore the two expression patterns cannot be compared.

To resolve this problem a simple approach would be to isolate barcoded identifiers from the human cluster, translate human gene measurements into homologous mouse genes, and re-assemble the expression data, similar to strategies used by others (Butler et al., 2018). This requires a unique (1:1) translation table to be constructed that reasonably compares expression patterns between the two species. Mouse-human homology tables generally have multiple matches that would need to be resolved or filtered. Furthermore, it is not clear if alignment of raw sequences to a combined reference genome would underestimate counted gene expression in the appropriate species due to some reads having inappropriate alignment with the wrong genome. One can devise several alternate strategies for translating and combining reads from multiple species— which is the most effective? To identify the optimal strategy, this study will evaluate differences among the methods, and compare methods for combining cells from multiple species. We will evaluate the specificity of species assignment of barcodes and we will identify gene expression differences between two species for the same clustered cell type (microglia).

## Results

### Transplants and sequencing

Human presumptive macrophage precursor (PMP) cells were prepared from iPSC and transplanted into the cortices of newborn (P0) immunocompromised (rag2^−/−^), hCSF-expressing mice as described (Xu et al., 2019). Four xenografted mice (two males and two females, labeled s1 through s4) were harvested 6 months after transplant. Isolation and dissociation of mouse brain tissue, single-cell isolation, and library preparation were described previously (Xu et al., 2019). Raw sequencing results are archived at NIH GEO, accession number <u>GSE129178</u>.

### Combined analysis

10X Genomics cellranger software was used to align raw sequencing reads (n=4 samples, mean 98,245,520 reads/sample) with a mixture of human (hg19) and mouse (mm10) reference genomes, resulting in an average of 112,048,732 alignments/sample. Barcoded counts per gene were loaded into the R package Seurat (Satija et al., 2015). After normalization, scaling, and clustering, one cluster was clearly enriched with human cells (**Figure 1A**, labeled as Human Microglia). A list of 1,684 barcodes identifying this human-specific cluster was exported for use in subsequent analyses. Another list of the remaining 17,459 barcodes was also exported, presumably corresponding to mouse cells.

**Figure 1.**
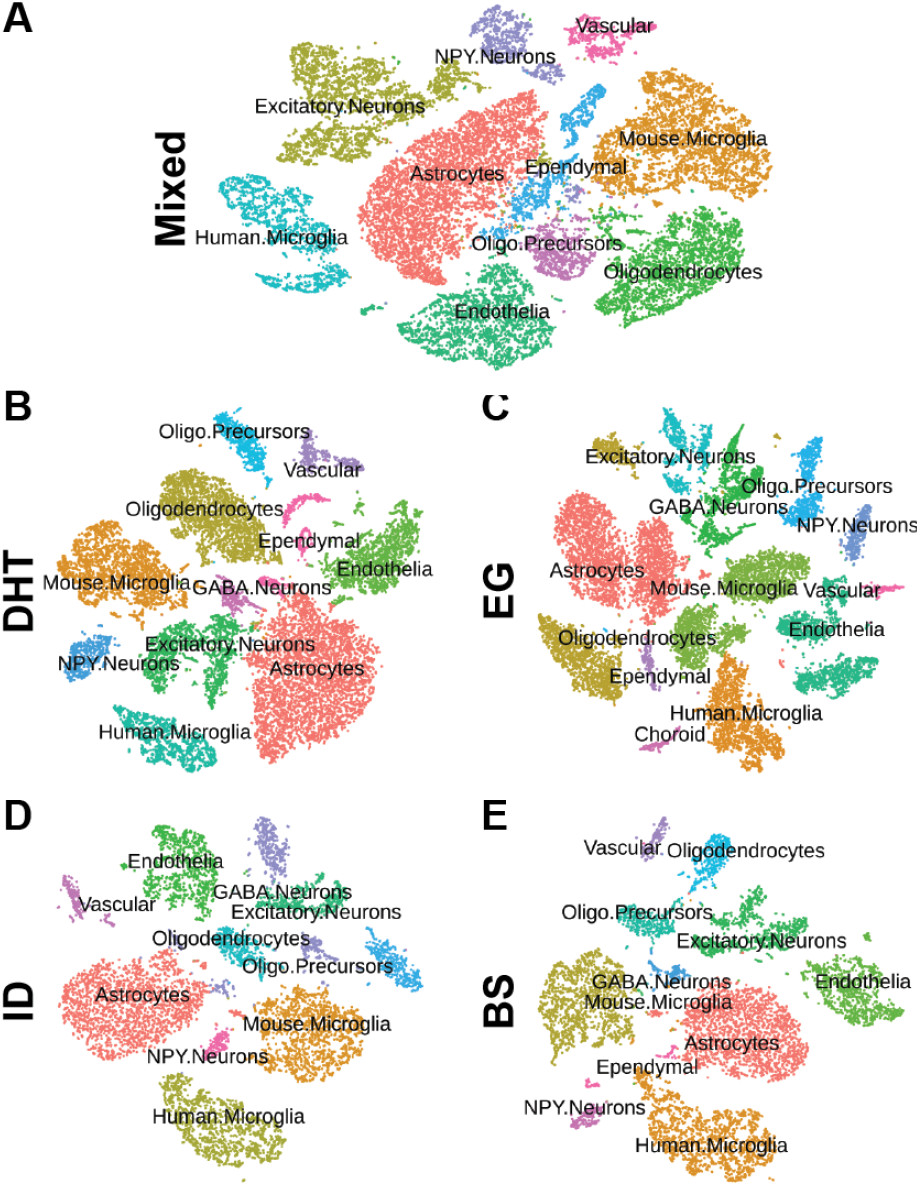
Clustering by Method. Individual cells were clustered and plotted as a tSNE projection. Cell types were labeled according to top genes differentiating each cluster from all others. (A) Mixed: All samples were aligned with a mixture of mouse and human reference genomes (mm10 and hg19). In this case the Human Microglial cluster primarily expresses human genes (gene symbol prepended with “hg19_”) and all other clusters primarily express mouse genes (symbol prepended with “mm10_”). (B) DHT: The human microglial clustered barcode identifiers were used to isolate a count table, which was translated to mouse symbols and merged with the remaining barcodes. (C) EG: The original FASTQ files were aligned with human and mouse reference genomes separately. The human count table was translated to mouse gene symbols and merged with the mouse count table. (D) ID: The Human Microglial cluster (from panel A) was used to extract sequencing read IDs from the mixed-genome BAM file. These were used to split the original FASTQ files into human and mouse-specific files, which were each aligned with the appropriate genome. The human counts table was translated to mouse symbols and merged with the mouse counts table. (E) BS: The mixed-genome BAM file was scanned to find the genome with the best alignment score for each read ID. These read IDs were used to split the original FASTQ files, which were then processed as in the ID method.

### Homologene table

Many human gene symbols correspond to multiple orthologous mouse genes, and vice versa. To obtain a table of unique gene symbols, a homologene table was downloaded from Jackson Labs (2019). This list was hand-curated to produce a unique list of 17,629 pairs of mouse-human matches (**<u>Supplemental File: geneTrans.txt</u>**). This list was edited by searching for duplicated gene symbols, removing symbols that were derived from clone names when traditional gene symbols were also available and choosing among numbered or lettered sets of symbol codes. For example, human *IL11RA* is homologous to mouse Il11ra1, Il11ra2 and Gm13305. Gm13305, which is derived from a name in a cDNA collection, designates a predicted gene overlapping Il11ra2. For this homology table, only Il1ra1 was retained and both Il11ra2 and Gm13305 were deleted. Similarly, if multiple gene orthologs were listed, and they were clearly labeled as an ordered series (e.g., with suffixes such as a,b,c…), only the first element was chosen. Any genes with multiple homologs that could not be resolved with these simple rules were deleted. The editing process, while somewhat arbitrary, was designed to produce a reasonable sampling of genes with known orthology between species. During analysis of scRNAseq data it was further determined that mitochondrial genes were missing from this list, and so they were added during sample processing (33 genes).

### Summary of Strategies

The initial approach was to extract human cell results from the genes × barcodes matrix, convert human gene symbols directly to mouse using the homologene table, and recombine results with mouse cells (this method is named Direct Homologene Translation [DHT]). A second strategy was to simply align the original FASTQ files with each genome separately (named Each Genome [EG]). For this to be effective, the 100 nt cDNA reads would need to be sufficiently distinct between species so as not to exacerbate the problem of inappropriate alignment. A third strategy would also rely on the human barcode list but, instead of re-naming existing gene counts, use it to split the original FASTQ reads into two species-specific subsets, which would each be aligned with only the appropriate genome (Split by barcode identifier [ID]). This would have the advantage of eliminating cross-species alignment but of course it would be limited by the accuracy of the initial clustering. Finally, the process of parsing and splitting read files could be extended to ignore the human-specific barcode list and instead directly split reads according to which genome produced the best alignment score for each sequencing read (Split by best score [BS]). The split FASTQ files could then be re-aligned with only the appropriate genome and re-assembled as before. Each of these will be evaluated by comparing the splitting of reads and barcodes into distinct genomes and how well each method drives the comparison of mouse and human microglia.

### Direct homologene translation (DHT)

For this strategy, sequencing counts were loaded from the mixed-genome cellranger count output into a Seurat data object with each replicate distinguished by prepending a sample identifier tag (s1…s4) to the barcode sequence tags. Raw counts were exported to a table in R. The human cells were selected using the barcodes exported from the initial combined analysis. In this subset table, only human genes appearing on the homologene table were preserved and translated from human to mouse gene symbol, resulting in a table of 9,689 gene symbols for 1,684 barcodes. The raw data table was similarly filtered using the mouse barcode array and only the mouse gene symbols present in the homologene table, producing a table of 14,828 gene symbols for 17,459 barcodes. These tables were re-loaded into separate Seurat data objects and then merged into a single object (15,022 genes × 19,143 barcodes). The R commands used for this process and diagnostic output are displayed in an RMarkdown report (**<u>Supplemental File 1)</u>**. The tSNE plot from this strategy (**Figure 1B**), shows a different pattern of clusters but overall a similar distribution of mouse brain cell types as the combined analysis (**Figure 1A**).

### Alignment with each genome (EG)

Raw FASTQ files were aligned separately to both human (hg19) and mouse (mm10) reference genomes using the cellranger count command, resulting in a mean 5,479,238 alignments per sample to hg19 and 103,991,443 alignments to mm10. Counts from each genome were loaded into distinct Seurat objects. The human object was exported to a table of raw counts, converted to mouse symbols using the homologene table, and re-imported into a Seurat object (11,377 gene symbols × 16,708 barcodes). The mouse object was selected to include only the mouse genes present in the homologene table (14,545 symbols × 29,055 barcodes). The two objects were then merged (14,781 gene symbols × 45,763 barcodes). An RMarkdown report for this procedure is shown in **<u>Supplemental File 2</u>**. A tSNE plot from this strategy (**Figure 1C**) splits several cell type clusters into distinct sub-clusters.

### Splitting sequencing reads by clustered barcode (ID)

With the list of human-specific barcodes from the initial clustering, we sought to select reads from the original FASTQ files that could be assigned as human sequences. Instead of beginning with the FASTQ file, this strategy required parsing the BAM file produced by cellranger count, since only this file contains the “corrected” barcode string that would perfectly match the human-specific list. A python script (<u>Supplemental File: splitByID.py</u>) iterated through the BAM file and stored FASTQ read IDs associated with the human barcodes from the cluster found in the combined analysis. The script then iterated through the paired FASTQ files, splitting them into two matching sets of barcode reads (R1) and cDNA sequences (R2) for each genome based on matching the selected read IDs. Each set was then separately aligned with the appropriate genome using cellranger. Counts were imported into Seurat objects. The human object was converted to mouse symbols using the homologene table (resulting in 9,336 gene symbols × 1,683 barcodes) and then merged with the mouse object, which was selected to include only genes matching the homologene table (14,526 gene symbols × 28,218 barcodes, resulting in a merged object of 14,725 gene symbols × 29,901 barcodes). An RMarkdown session using this procedure is found in **<u>Supplemental File 3</u>**. Results (**Figure 1D**) produce a more uniformly coalesced set of clusters, with human and mouse microglia adjacent on the plot.

### Splitting raw sequencing reads by best alignment score (BS)

During preparation of the ID python script, we realized that the BAM file also contained the alignment score for each read matching one or both genomes. We reasoned that alignment with the correct species would produce the better alignment score. The python script was revised to iterate through the BAM file, store the read ID, the alignment score and the aligned species, retaining only the one with the highest alignment score for each read ID. The original list of barcode IDs was not used. The split FASTQ files were aligned with the appropriate genome using cellranger. As before, counts from each set of alignments were loaded into Seurat, the human matrix was translated to mouse symbols (9,337 gene symbols × 3,354 barcodes), and combined with the mouse matrix containing only homologous gene symbols (14,523 gene symbols × 28,237 barcodes; resulting in an object with 14,745 gene symbols × 31,591 barcodes). The RMarkdown session for this procedure is found in <u>Supplemental File 4</u> and the python script can be viewed on a GitHub repository file (<u>Supplementary File: splitByScore.py</u>). Results (**Figure 1E**) exhibit distinct clusters with greater separation.

### Splitting FASTQ files

Two of the strategies split the original reads into two species-specific sets of FASTQ files for re-alignment (**Figure 2**). Using only the selected human cluster barcode IDs from the initial analysis, after splitting, the human-specific reads averaged 5,469,135 per sample (5.6% of total) and the mouse reads averaged 92,776,385 per sample (94.4%). Splitting by best alignment score slightly increased the number of detected human reads to an average of 8,233,854 (8.4%) with lower mouse reads (83,422,959; 84.9%). In one example, a read having an alignment score of 99 on human chr10 was not part of the barcode list used to split by ID, so in the ID strategy it aligned to mouse chr1 with alignment score of 35. The ID method, then, counted the read towards an incorrect gene while the BS method correctly partitioned the read to the human genome. While this is only one limited example, this illustrates that it is possible for a low-quality alignment to be split to the wrong genome and mis-counted when a less accurate strategy is employed, such as ID.

**Figure 2.**
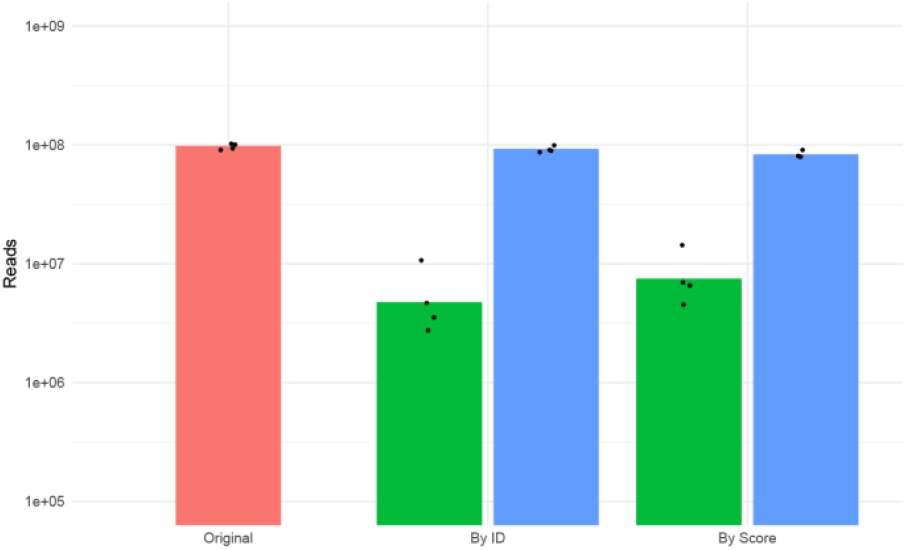
Numbers of reads aligned to each genome. The bars show the mean numbers of reads aligned with each genome (red: mixed; green: human; blue: mouse), by each strategy, plotted on a log scale. Dots show the numbers of reads for each sample (n=4).

The numbers of aligned reads largely reflected the split of FASTQ reads for the ID and BS methods, but were surprisingly nonspecific for the EG method, which did not split the FASTQ files. Over 20% of the number of total alignments in mixed genomes were identified in human genome, much more than any other method. This likely demonstrates that an acceptable alignment can be found for similar exon sequences in both genomes.

### Separation of Barcodes by Species

After loading all recombined tables into Seurat objects, the numbers of barcodes identified by species for each method were compared for genome selectivity and overlap (**Figure 3**). In this analysis we counted only cells with at least 500 genes detected above 0 counts per gene. Not surprisingly based on the original algorithm, the DHT method had no overlap in barcodes because one cluster of barcodes was used to split the table of counts by species (**Figure 3A**). Similarly, using the same cluster of barcodes to split the original reads into two species-specific files (ID method, **Figure 3C**) also produced no overlap. However, slightly more human-specific barcodes were identified by ID than DHT methods (1,300 vs. 1,269, an increase of 2.4% for the ID method). This difference may be minor, but it exemplifies that DHT likely underestimates not only barcodes but also gene counts due to the presence of the alternate reference genome during alignment. The other two methods produced overlapping barcodes assigned to both genomes. The EG method had 1,001 overlapping barcodes (**Figure 3B**), which was judged to be unacceptably large and likely indicates the unexpected proportion of reads that aligned to both genomes. Splitting by score (BS, **Figure 3D**) produced only 147 overlapping barcodes, but also had a larger number of human-only barcodes (1,314).

**Figure 3.**
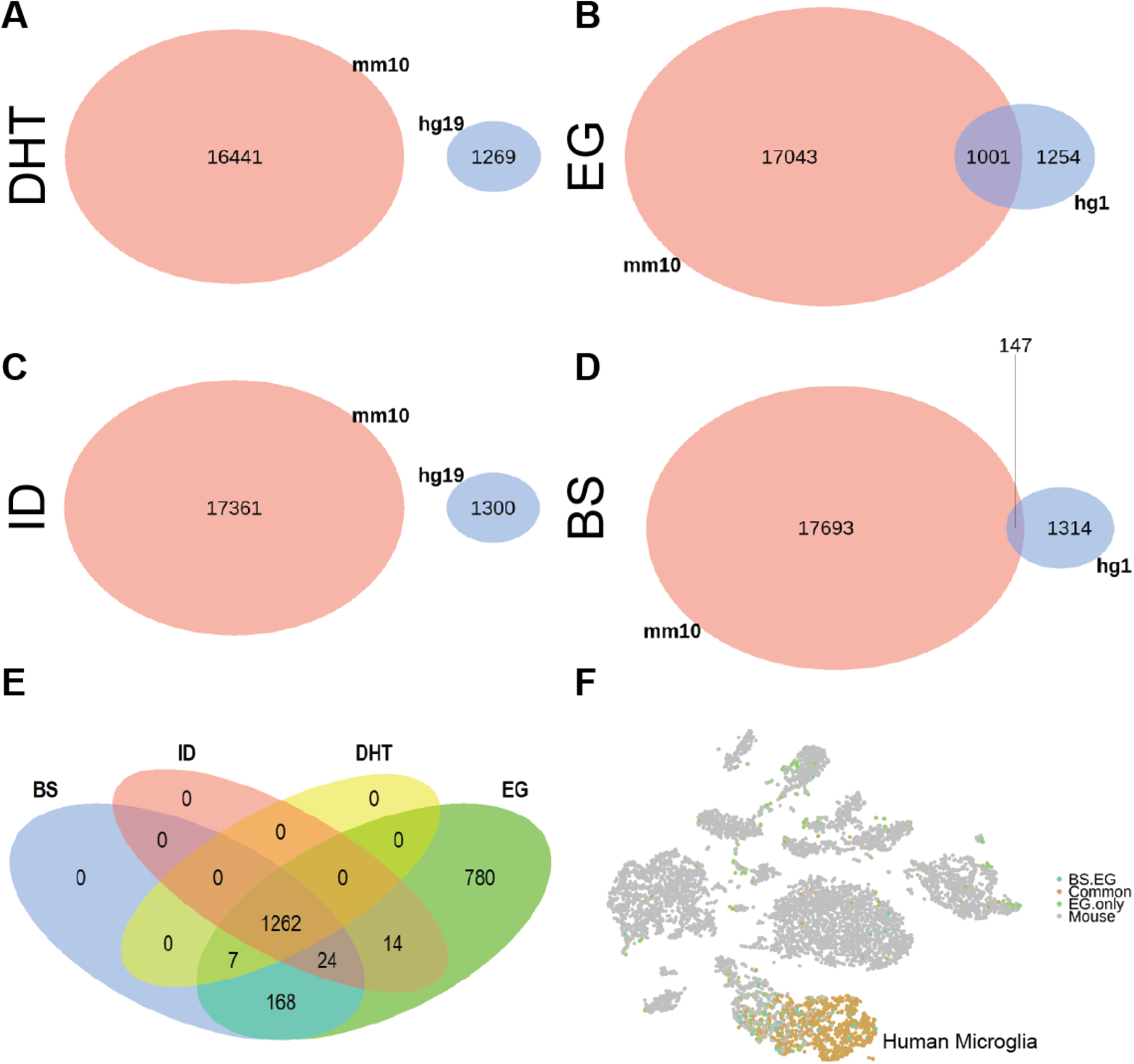
Separation of barcodes by species. (A-D) Venn diagrams of overlapping sample/barcode identifiers assigned to each species. For each method, the total numbers of sample/barcode identifiers (used as a proxy for numbers of cells) is shown for mouse (coral), human (light blue) and those found in both (shaded). Methods that distinctly separated counts (DHT) or reads (ID) showed no overlap. (A) DHT. (B) EG (C) ID (D) BS. (E) Human-assigned barcodes are compared for each method. (F) The BS tSNE plot (Figure 1E) was colored according to overlapping region from the Venn diagram (panel E). All barcodes assigned to human by all methods (Common, light orange) as well as those only identified as human by both BS and EG (BS.EG; blue-green) are found within the cluster of human microglia. Many of the barcodes assigned to human by EG (green) are scattered throughout the mouse cell types.

The barcodes assigned to human were compared for overlap and differences in a Venn diagram (**Figure 3E**). As expected EG identified a large number of barcodes (780) that were distinct from all other methods. Surprisingly, DHT identified only 7 barcodes overlapping with EG and the remainder (1,262) were common to all methods. 168 barcodes overlapped between BS and EG, but not ID or DHT, likely identifying human cells missing from the mixed-genome combined alignment (**Figure 1A**). To visualize the specificity of these subsets of barcodes, the tSNE plot (as calculated with the BS method, **Figure 1E**) was color-coded by the major groups from the Venn diagram (**Figure 3F**). Both the Common (1,262 barcodes common to all methods; light orange) and the overlap between EG and BS (168 barcodes; blue) overlaid the human microglia cluster nicely, but the EG-specific barcodes (780; green) were spread throughout the plot, seemingly a random subset of all cells. These comparisons indicate that while a core group of barcodes could be identified by any of the four methods, the EG method produced barcodes inconsistent with the one human cell type (microglia) and therefore this strategy should be avoided. Only the BS method identified a larger number of human cells with little evidence of misidentifying mouse cells.

To better compare the causes of multi-species overlaps, 15 randomly-selected barcodes were chosen from the 147 overlapping by BS (**Figure 3D**). Summarizing the distribution of alignment scores per read by genome (**Supplemental Figure 1**), as determined from BS, EG, or ID bam files, there was clear indication that these barcodes likely represented two-cell droplets. That is, reads encoded with each of the 15 sampled barcodes exhibited a high distribution of alignment scores in both species for either BS or EG methods. As expected, some barcodes were not members of the combined-genome human barcode list and therefore had no results from the ID method. This conclusion was confirmed by determining that there was a similar number of reads in each genome per barcode (**Supplemental Figure 2**). A direct comparison of alignment scores for each read from barcodes overlapping both genomes using the EG method (**Supplemental Figure 3**) showed that while most reads (here depicted as single dots) exhibited stronger alignment scores in one genome (dots plotted near either y=100 or x=100), most reads also produced relatively strong alignments in the alternate genome (dots between 70 and 100 in the alternative genome). That is, most reads in this sample exhibited strong alignment scores against both genomes. Therefore, this set of 147 barcodes should be removed from the analysis as technical contaminants since they were likely produced by adding more than one cell to the microfluidic droplet. Certainly, this indicates that other double-cell barcodes also exist in the dataset but would be undetectable since, in most cases, both cells would likely derive from the same species. However, only 147 out of 19,154 cells, or 0.76%, could be identified as two-cell droplets, suggesting that the scale of this problem is relatively small.

In contrast, a randomly-selected set of 15 barcodes in the EG overlap that were absent in the BS overlap produced a different pattern of results (**Supplemental Figures 4-6**). In several cases there were much larger discrepancies in summarized alignment scores by genome, read counts, and individual read alignment scores across both species. In this set most barcodes were enriched in mouse-specific alignments but with lower-quality alignments to human genome. Finding evidence of reads that align with both species in this analysis demonstrates why using any method that allows individual reads to align with more than one species (DHT or EG) should be unacceptable.

### Comparison of mouse vs. human microglia gene expression

A biologically-relevant measurement to compare the methods is to use each expression dataset to find significantly different genes between converted human and mouse microglia. This was a major goal of our initial analysis (using the DHT algorithm), where we found evidence of human-specific expression differences in human microglia, including several classes of disease-associated genes (Xu et al., 2019). To compare methods, lists of human vs. mouse microglia genes found by Seurat using the FindMarkers function by contrasting human-specific and mouse-specific clusters identified by expression of the characteristic human and mouse-specific genes, Spp1 and Hexb, respectively. The p-value for difference was calculated by a Wilcoxon Rank Sum test with a default log fold change of 0.25 and minimum percent expression of 0.1, with a Bonferroni correction applied. For functional analysis, a stringent filter was applied, requiring an adjusted p-value of 0.05 or less and a log fold change of 1 or more (2-fold). Using these methods, the DHT algorithm found 1,711 genes with 246 passing the stringent filter. Other methods identified: EG, 1,258 total with 161 passing the stringent filter; ID, 1,728 total with 234 stringent; and BS, 1,483 total with 177 passing the stringent filter. These lists were compared by Venn diagram on the total differences (**Figure 4A**) or the stringent differences (**Figure 4B**). In each case there was a common core of genes selected by all methods (917 for the total lists, 121 for the stringent lists). BS appeared to exhibit the fewest differences from the common list among the stringent list (56 additional genes, of which only 2 were not in common with at least one other method).

**Figure 4.**
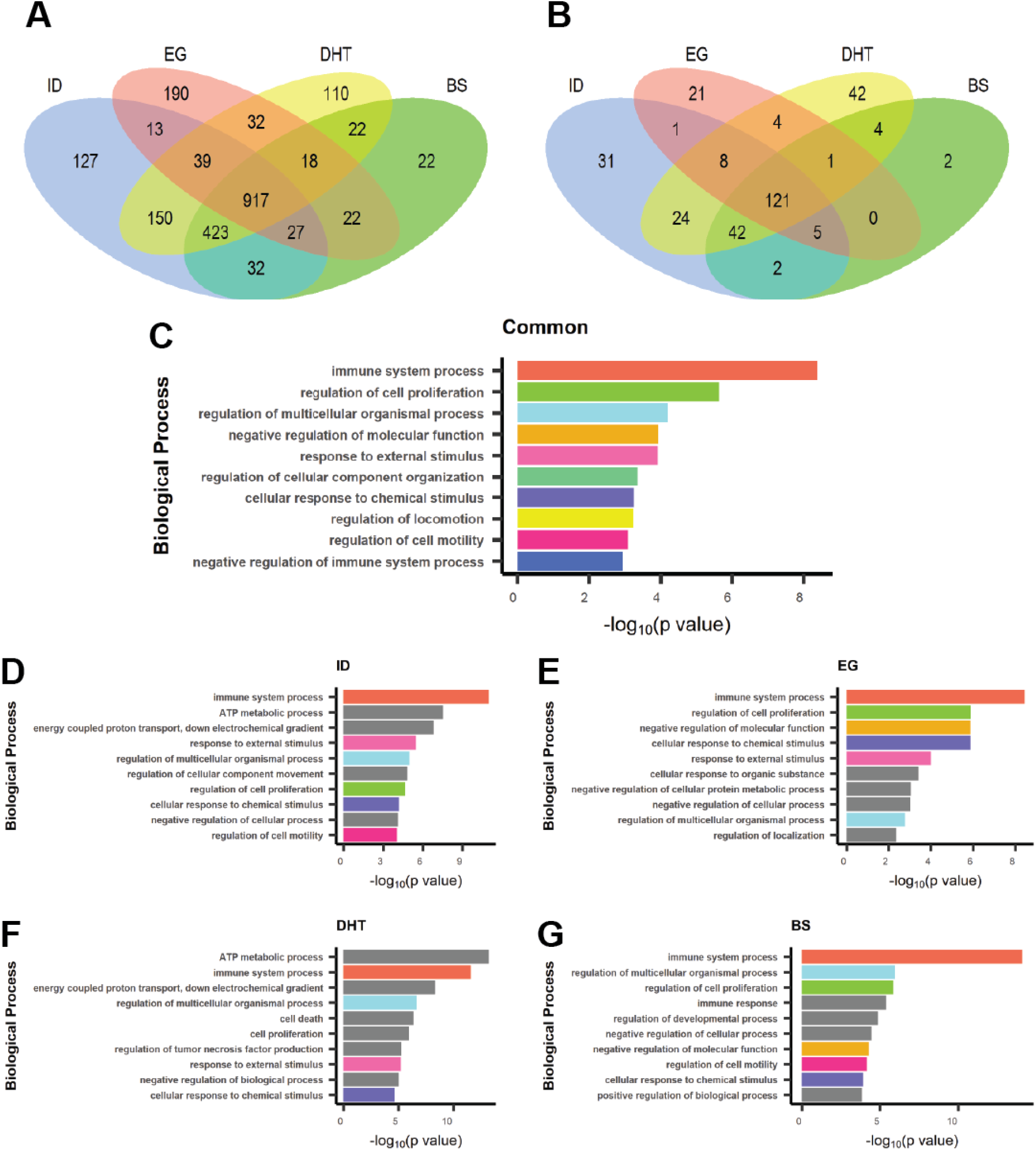
Gene expression differences between mouse and human microglia. Following clustering and cell type assignment, genes differing between mouse and human microglial clusters were obtained. (A) Overlap of all significantly different genes. (B) Overlap of high-stringency gene differences (p≤0.05 and a log2 fold-change of 1 or greater). (C) The 121 high-stringency genes found in common to all methods were used to identify enriched gene ontology biological (GO-BP) process terms. The top 10 terms, by minimal p-value, are shown. (D-G) Top 10 GO-BP enriched terms for each method, with bars colored if they match a top 10 term from the common gene analyses in panel C.

Taking the common genes from the stringent lists (**Figure 4B**, 121 overlap), functional analysis was run with g:Profiler (Reimand et al., 2016) focusing only on biological processes (**Figure 4C**). The top 10 (by smallest p-values) included many expected functional terms relating to microglia including immune system process, response to external stimuli, and cellular response to chemical stimulus. All of these would be expected to be enriched in microglia, although it is interesting that these are differences between mouse and human microglia. There are several studies demonstrating that mouse and human microglia are distinct with characteristic gene expression patterns (Gosselin et al., 2017; Galatro et al., 2017; Xu et al., 2019). When stringent differential expression lists of genes from each method were similarly analyzed for enriched biological processes, there was substantial overlap with the common list but also many differences (**Figure 4D-G**). For comparison, the colors for each GO term in the common list (**Figure 4C**) were reproduced in each method-specific barplot and those not found in the common list are colored as gray. Again, several functional terms expected to be associated with microglia can be found among the method-specific lists. The BS plot (**Figure 4G**), for example, has an additional immune response term as well as several regulatory terms. The differences among these functional summaries exemplifies why the choice of algorithm is important for biological interpretation. However, there is no clear conclusion whether any of these is more appropriate or correct according to the biology of mouse vs. human microglia.

### Optimized by-score method (BS2)

Since the partitioning of reads into appropriate genome according to the best alignment score appears to be the most appropriate strategy, and since this method reveals a small number of barcode-tagged “cells” that likely represent mixtures of mouse and human cells, we concluded that the optimal strategy was BS with exclusion of overlapping barcodes. Therefore, we ran the BS algorithm but during the process of converting human to mouse identifiers, we also excluded 412 barcodes (columns in the tables) that were common to both the mouse and human datasets. The resulting human table included 9,337 gene symbols × 2,942 barcodes. The mouse table included 14,523 gene symbols × 27,825 barcodes. The combined Seurat object included 14,725 gene symbols × 30,767 barcodes. After scaling, dimension reduction, and clustering, a slightly clearer tSNE plot emerges (**Figure 5A**). The RMarkdown session for this procedure is found in <u>Supplemental File 5.</u> There are no overlapping barcodes (**Figure 5B**), as expected. This revised analysis produced a clear set of enriched gene ontology/biological process terms (**Figure 5C**), that were further split into the human-enriched terms (**Figure 5D**) and the mouse-enriched terms (**Figure 5E**). Here we find that immune system response is present in both species of microglia, but with different enriched sets of individual genes. Prominent human-selective terms include regulation of cytokine production, regulation of cell proliferation, cell death, and T cell activation. Mouse-selective terms include regulation of cell motility and developmental processes.

**Figure 5.**
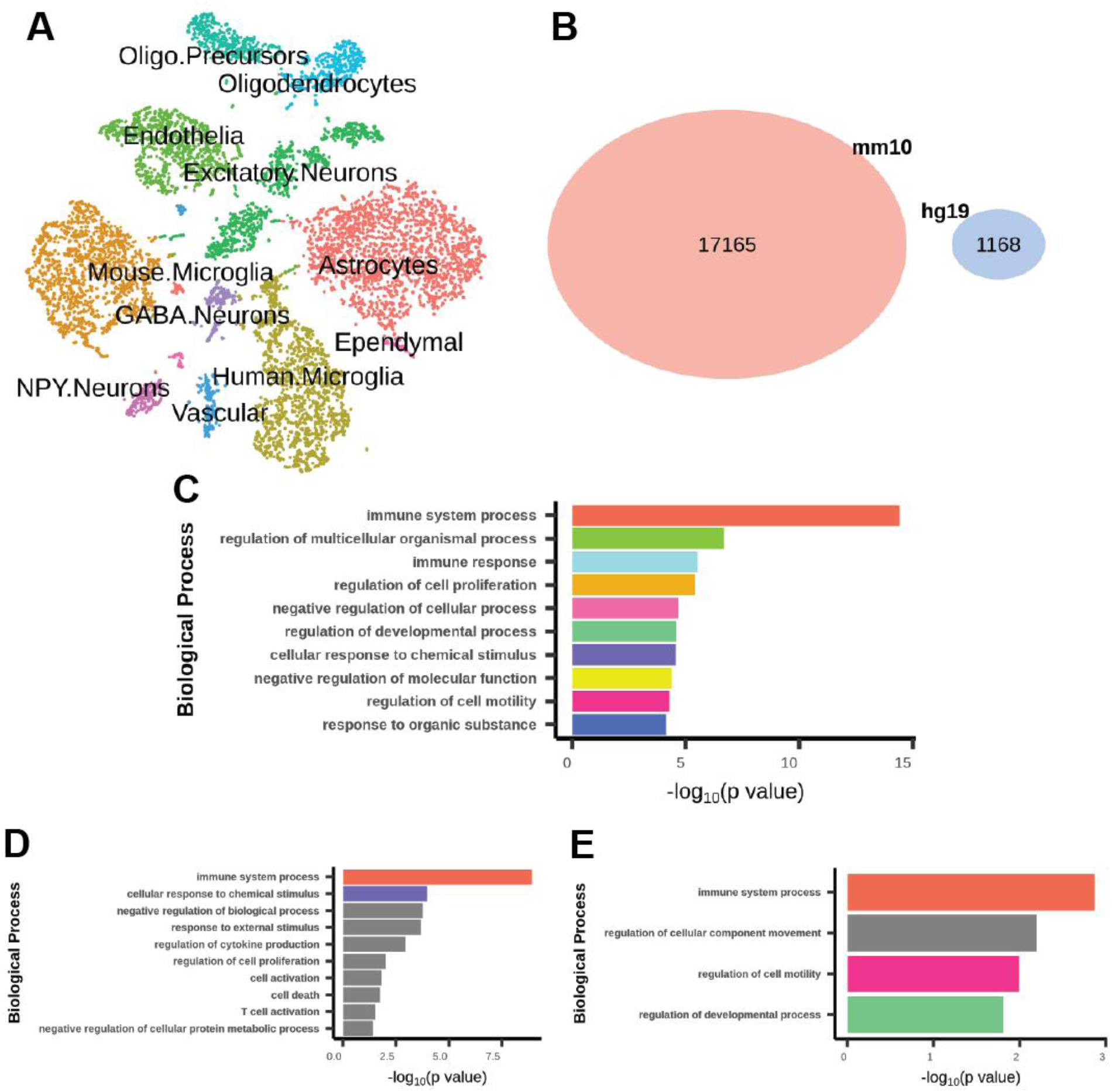
Optimized method: Split by score excluding overlaps. The BP algorithm was run as before except that barcodes found in both species were excluded after gene translation but prior to merging species. (A) tSNE plot labeled by cell type. (B) Number of barcodes assigned to each genome, with no overlap by design. (C) Top 10 GO-BP terms from genes different between the human and mouse microglial clusters. (D) Top 10 GO-BP terms enriched in the human cluster. (E) Top 4 GO-BP terms enriched in the mouse cluster (only 4 terms passed the p-value threshold). Bar colors match terms shown in panel C.

## Conclusions

The goal was to devise and compare several methods for partitioning scRNAseq data into two species. Analysis of the results guides reasonable choices for optimal strategies. Clearly, while mixed-species scRNAseq results may be initially evaluated by aligning with mixed genomes, this is not an appropriate way to perform final analyses since there is evidence of sequencing reads aligning with reasonable quality to both genomes. Alternatively, reads from a single barcode (assumed to be a single cell) may optimally align with two different species, and in some cases this may be due to droplets containing multiple cells (for example, **Supplemental Figure 6**). Many barcodes overlapping both species were identified with this characteristic, and the majority of these have better alignments to one of the species (**Figure 3B** [EG] and **Supplemental Figures 4-6**). The recommendation is, then, to use a pre-processing strategy for parsing FASTQ files of raw reads into the appropriate species before analyzing gene expression.

We evaluated strategies to use initial, mixed-genome clustering to split FASTQ files (ID) or to split by best alignment score (BS). As an extreme case for comparison, we aligned all reads to each genome independently (EG). The latter approach is clearly the most problematic, with the largest number of barcodes assigned to both species (**Figure 3**). However, splitting by ID incorporated mixed-species barcodes, as outlined above, and excluded 168 shared by BS and EG (**Figure 3E**) that clustered with human microglial cells (**Figure 3F**). EG also produced an unacceptable number of human-selected barcodes clustering with non-microglial cells (**Figure 3F**). Neither EG nor ID was judged to be ideal. Similarly, DHT had slightly fewer barcodes assigned to human and there was concern that some genes might be undercounted or miscounted due to the presence of the alternate genome during alignment. This argues in favor of splitting the sequencing reads before final alignment with a single reference genome.

Among methods tested for splitting sequencing reads by species, it was found that the optimal method was to split by alignment score. This produced the fewest number of barcodes that clustered with cells of the alternate species (**Figure 3F**). However, since a small number of barcodes parsed to both genomes (147 barcodes in **Figure 3D**), we extended the BS algorithm to exclude any overlapping barcodes. In this revised BS2 method, 412 barcodes were deleted, likely leading to the 147 from Figure 3D after filtering and scaling of the data. The BS2 approach produced distinct tSNE clusters (**Figure 5A**) with no overlap among barcodes (**Figure 5B**).

Differential expression analysis contrasting human and mouse microglia were largely consistent by method (**Figure 4**). However, distinct biological process terms were enriched when analyzing each method individually (**Figure 4D-G**). This exemplifies that the choice of method impacts functional biological interpretation. The final, revised BS2 approach gave a list of top ten enriched biological processes (**Figure 5C**) that was quite similar to the BS method (**Figure 4G**). Ideally, the removal of barcodes overlapping by species should have had little effect on the expression difference list.

We conclude that the optimal method for analyzing scRNAseq data from cells of multiple species is to pre-process the FASTQ files and split them by the best alignment score to each species. This requires scanning the BAM file produced from a mixed-species alignment, since this contains alignment scores that can be associated with sequencing read identifiers. With lists of species-specific identifiers, the original FASTQ files can be split into two species-specific sets. The now species-specific FASTQ files are then aligned only with the appropriate species, eliminating the major source of crossreactivity. Finally, barcodes found to be matched to both genomes are likely to arise from multiple-cell droplets in the partitioning technique and these should be excluded. To directly compare gene expression patterns, a unique mouse-human translation table was constructed that allowed merging of the two species-specific expression tables prior to analysis. Results indicate that this optimal method produces clear clustering by cell type and species, as well as identifying species-specific gene expression patterns in the same cell type (microglia). This approach should be valuable for all multiple-species scRNAseq experiments.

## Supplemental Materials

Sequencing data used for this study are available from the NIH GEO archive under accession number <u>GSE129178.</u> All supporting files are found on the “Mousify” project GitHub repository: https://github.com/rhart604/mousify. Note that the GitHub tool to serve html files does not display the paged table results used in the RMarkdown sessions. To view the complete file, download the html file from the GitHub project and open it with a web browser.

## Acknowledgements

This work was supported in part by grants from the NIH (R21HD091512 and R01NS102382 to Peng Jiang) R.P.H. was supported by R01ES026057, R01AA023797, and U10AA008401.

## Supplemental Figure Legends

**Supplemental Figure S1**. Sample alignment score summaries for barcodes overlapping human and mouse in the BS method. A random sample of 15 barcodes were selected from the 147 assigned to both human and mouse in Figure 3D by the BS method. Alignment scores were collected from the BS, EG, or ID methods for separated human (hg19) or mouse (mm10) genomes for sequencing reads with the selected barcodes. Alignment scores were summarized with a box and whiskers plot, with the central horizonal line representing the median, the top and bottom of the box as the interquartile range, and the whiskers showing the 95% confidence interval of the score distributions.

**Supplemental Figure S2**. Sample numbers of sequencing reads for barcodes overlapping human and mouse in the BS method. From the same data depicted in Figure S1, the total numbers of sequencing reads per genome per method are plotted as bars.

**Supplemental Figure S3**. Individual sequencing read alignment scores for barcodes overlapping human and mouse in the BS method. For each sequencing read (depicted as a dot), the alignment score is shown for the mouse genome (x axis, mm10) versus the human genome (y axis, hg19). Dots are only from the BS method.

**Supplemental Figure S4**. Sample alignment score summaries for barcodes overlapping human and mouse in the EG method but not the BS method. A random sample of barcodes was chosen from the 1,001 shown in Figure 3B that were not found in the 147 overlapping barcode from Figure 3D. Alignment scores were collected from the BS, EG, or ID methods for separated human (hg19) or mouse (mm10) genomes for sequencing reads with the selected barcodes. Alignment scores were summarized with a box and whiskers plot, with the central horizonal line representing the median, the top and bottom of the box as the interquartile range, and the whiskers showing the 95% confidence interval of the score distributions.

**Supplemental Figure S5**. Sample numbers of sequencing reads for barcodes overlapping human and mouse in the EG method but not the BS method. From the same data depicted in Figure S4, the total numbers of sequencing reads per genome per method are plotted as bars.

**Supplemental Figure S6**. Individual sequencing read alignment scores for barcodes overlapping human and mouse in the EG method but not the BS method. For each sequencing read (depicted as a dot), the alignment score is shown for the mouse genome (x axis, mm10) versus the human genome (y axis, hg19). Dots are only from the EG method.

