## Supplementary figures and images for "Strategies for Integrating Single-Cell RNA Sequencing Results With Multiple Species"

### Supplemental Figure 1

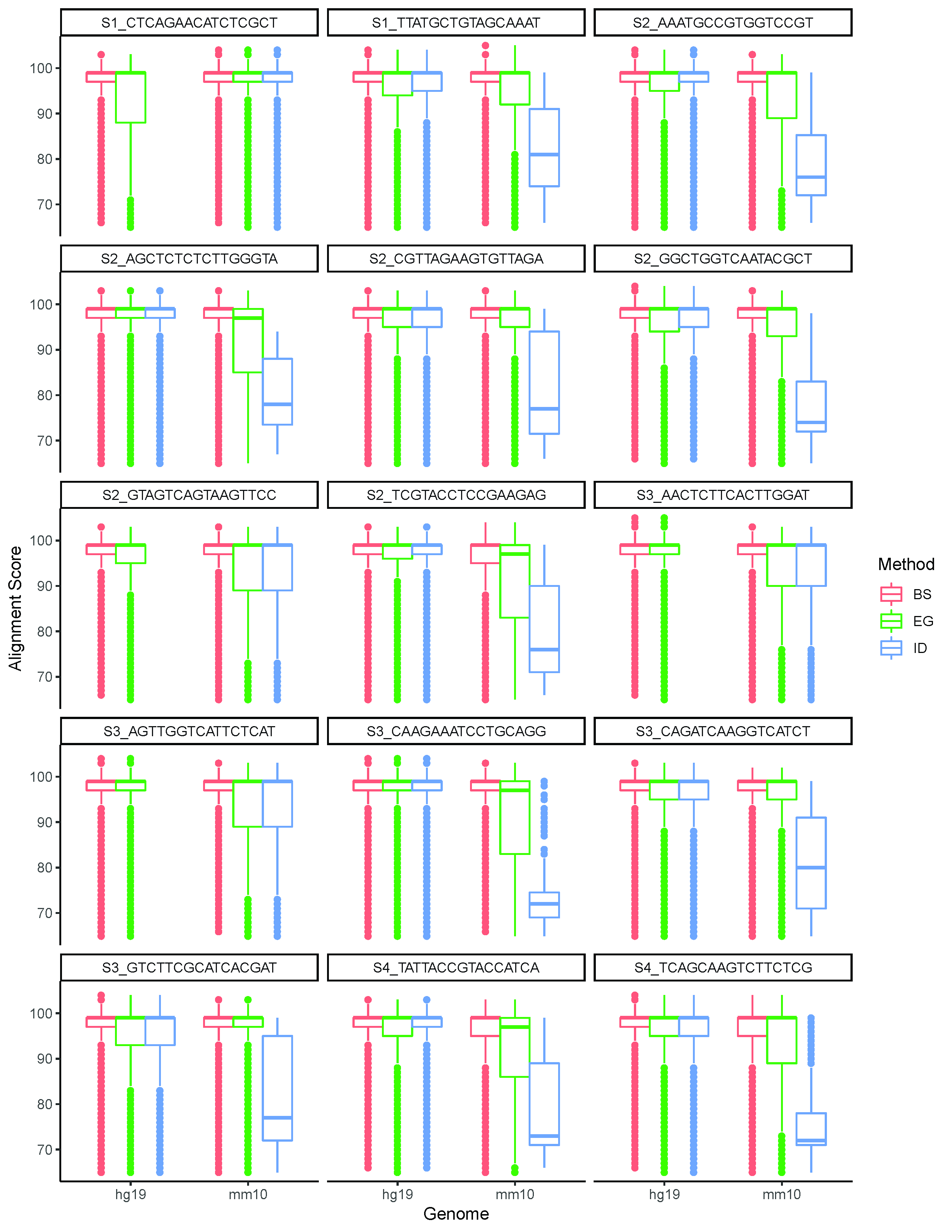

### Supplemental Figure 2

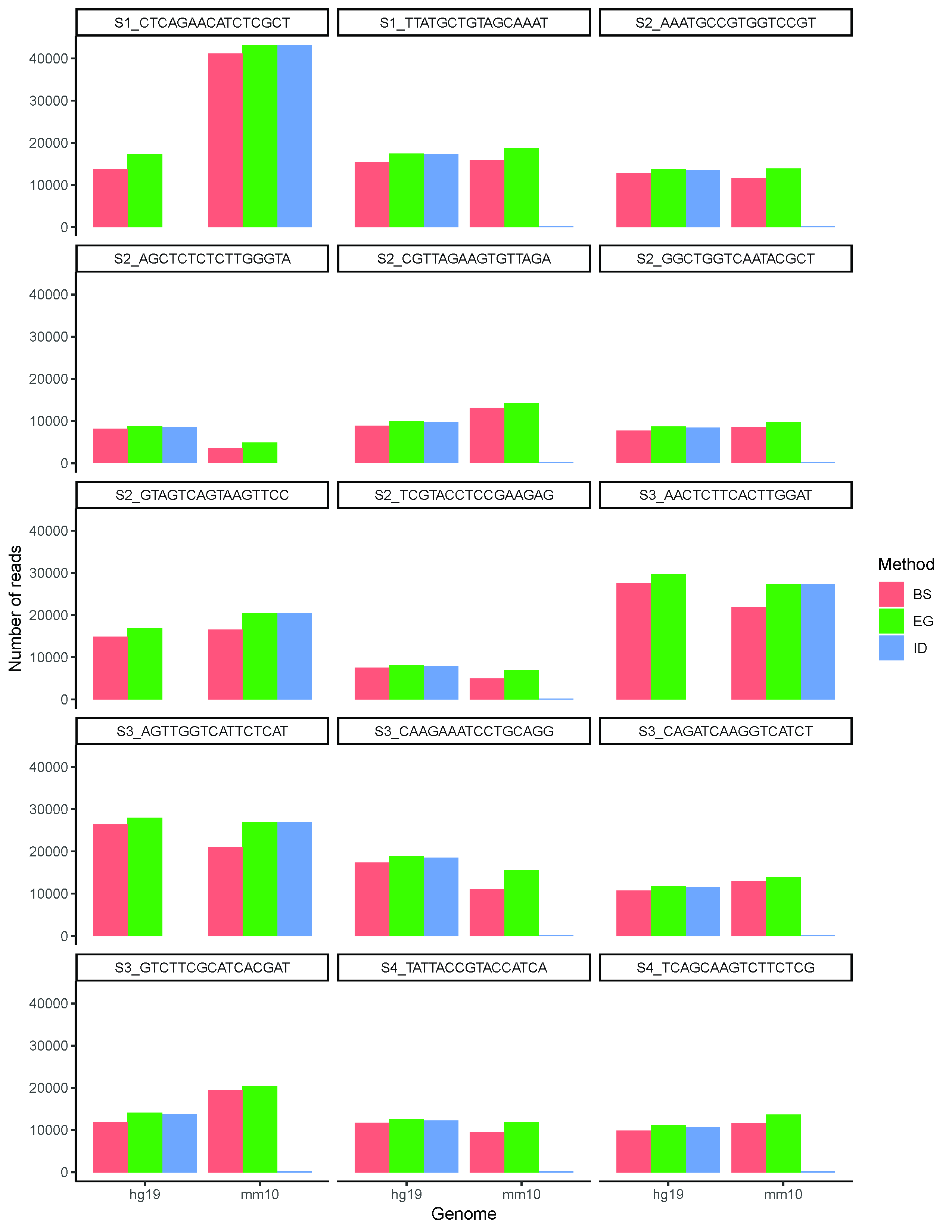

### Supplemental Figure 3

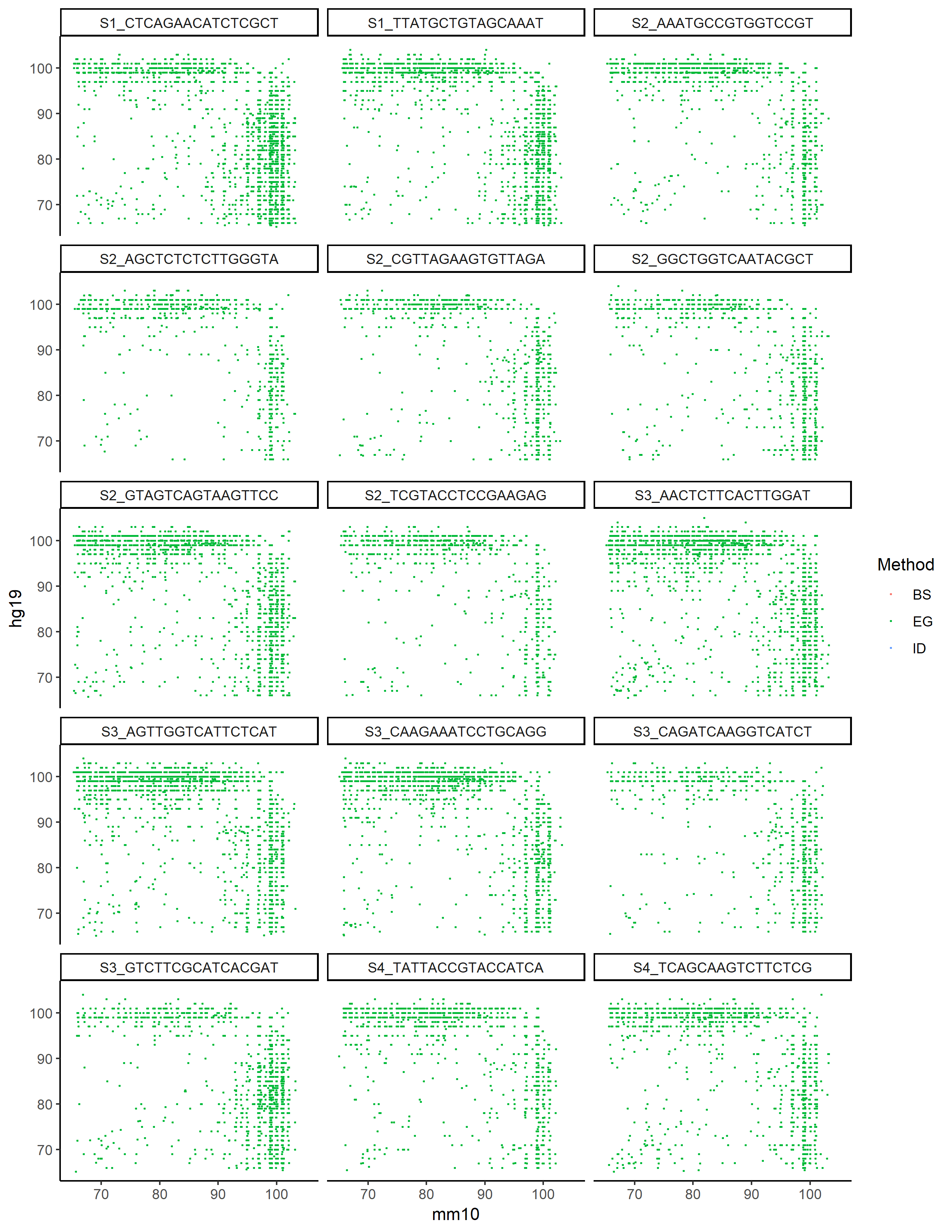

### Supplemental Figure 4

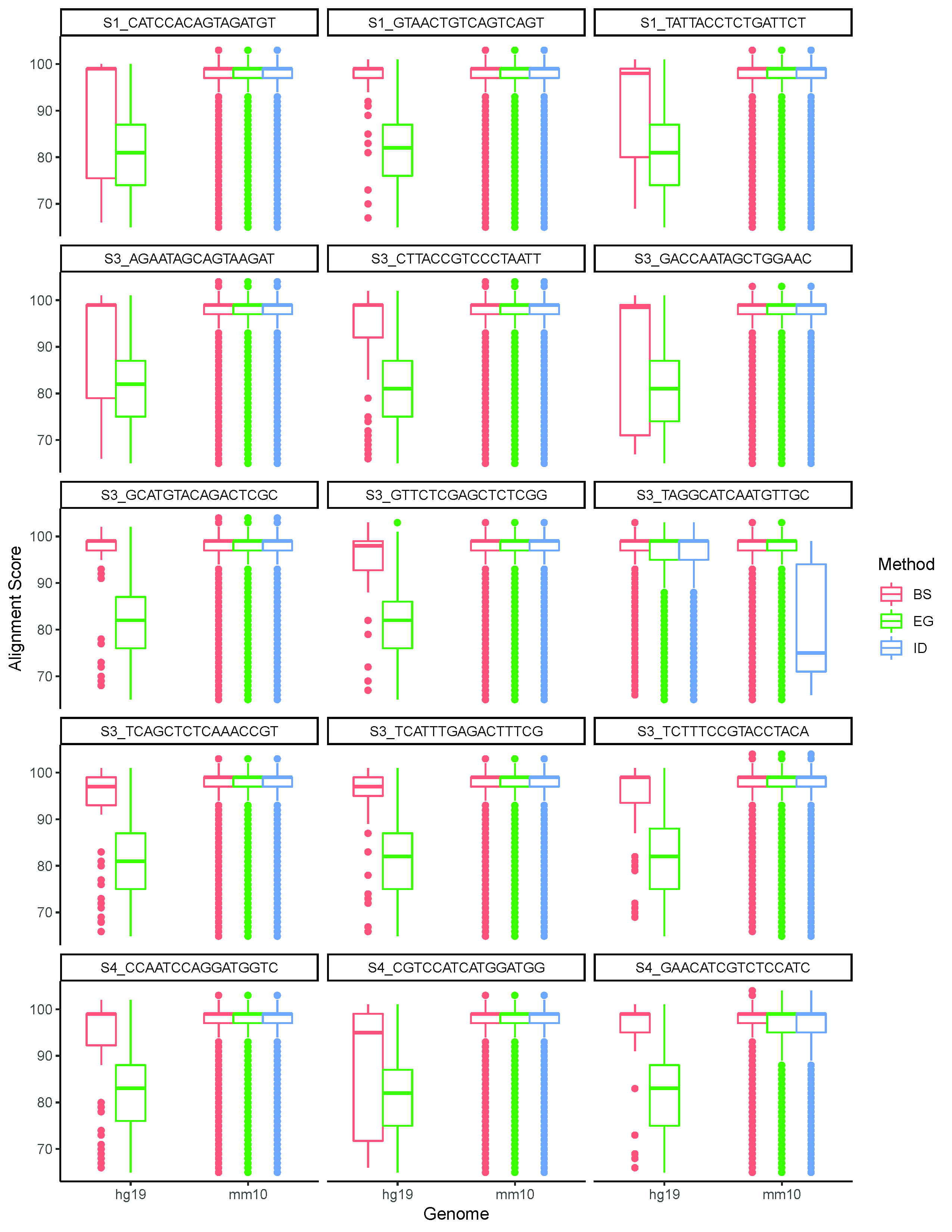

### Supplemental Figure 5

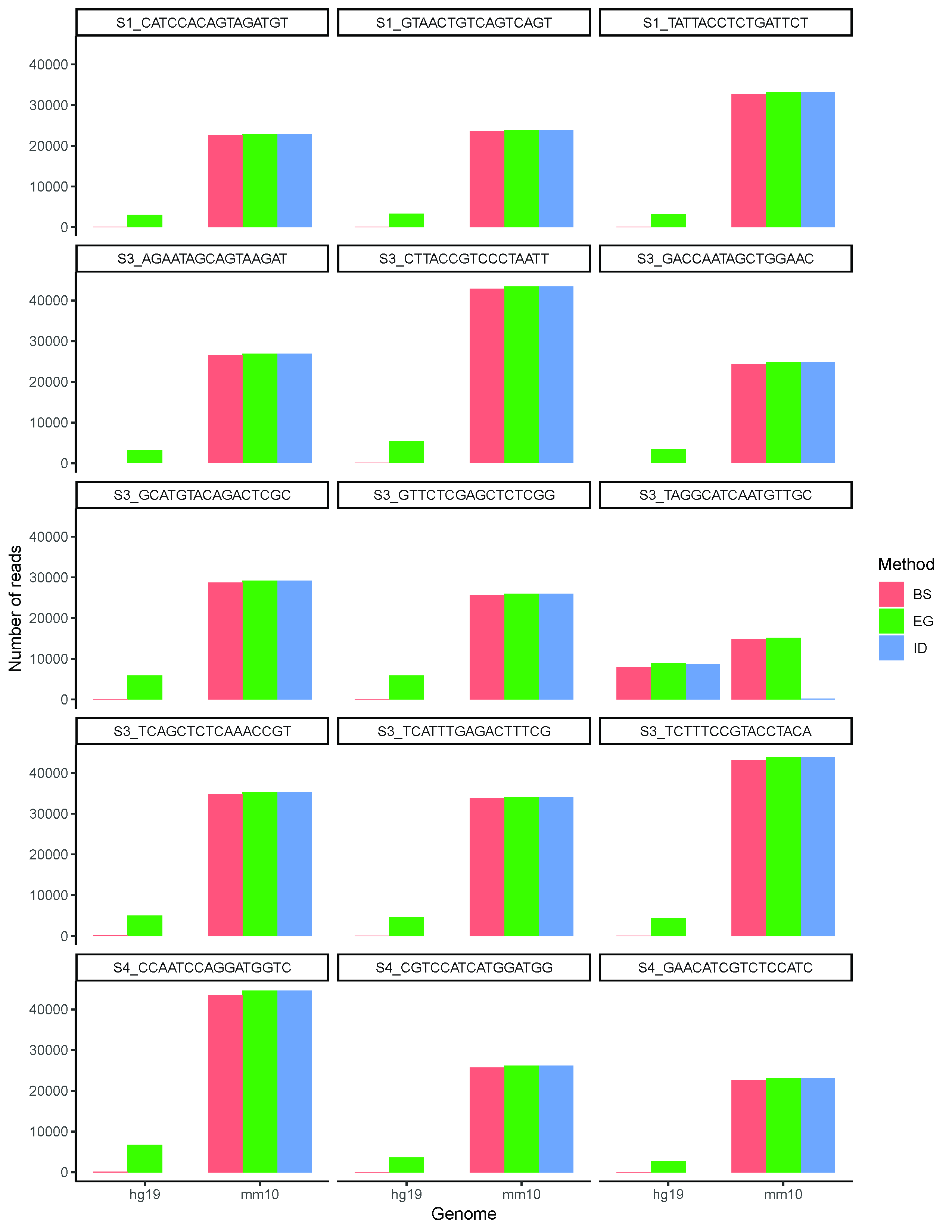

### Supplemental Figure 6

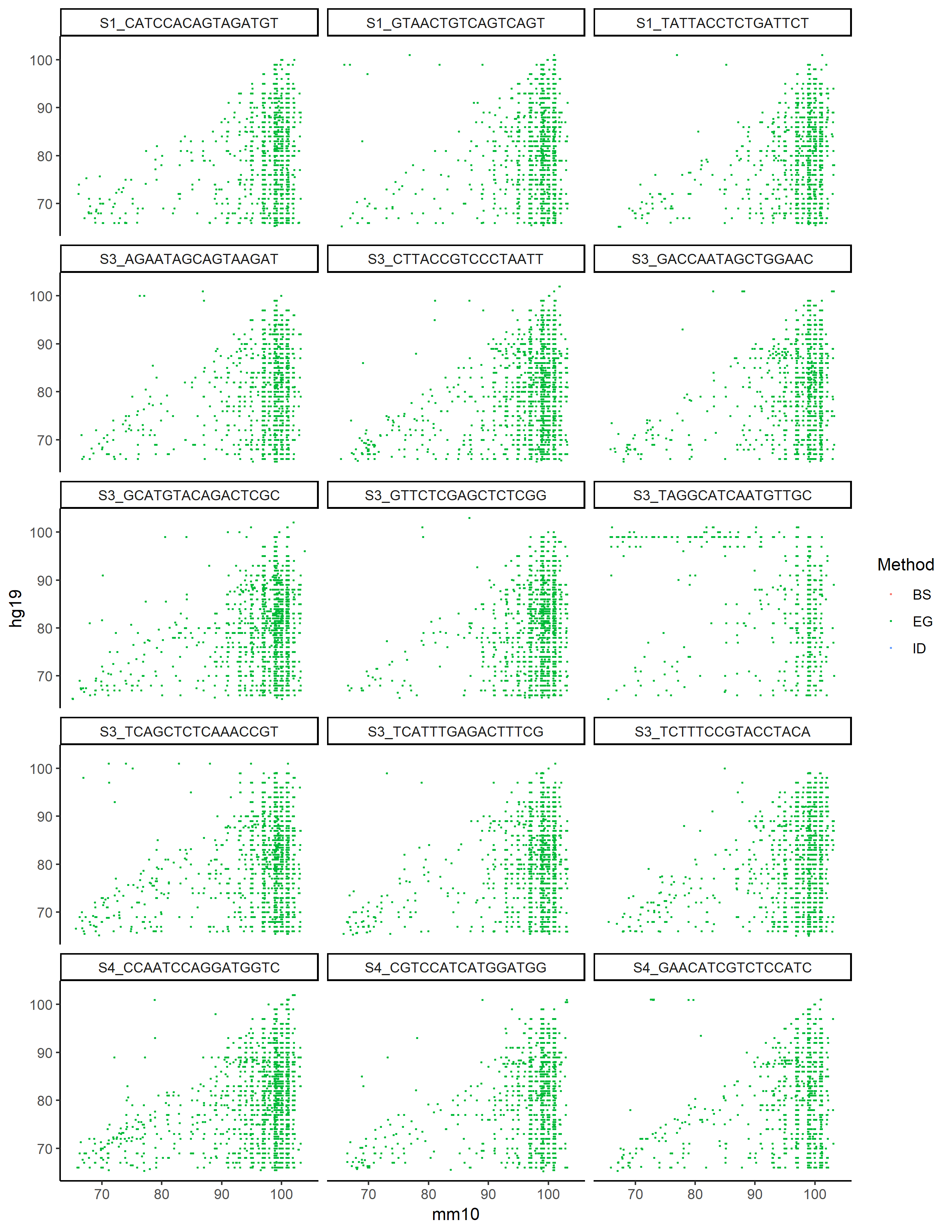
